# Lithium ions display weak interaction with amyloid-beta (Aβ) peptides and have minor effects on their aggregation

**DOI:** 10.1101/2021.01.03.425155

**Authors:** Elina Berntsson, Suman Paul, Faraz Vosough, Sabrina B. Sholts, Jüri Jarvet, Per M. Roos, Andreas Barth, Astrid Gräslund, Sebastian K. T. S. Wärmländer

## Abstract

Alzheimer’s disease (AD) is an incurable disease and the main cause of age-related dementia worldwide, despite decades of research. Treatment of AD with lithium (Li) has showed promising results, but the underlying mechanism is unclear. The pathological hallmark of AD brains is deposition of amyloid plaques, consisting mainly of amyloid-β (Aβ) peptides aggregated into amyloid fibrils. The plaques contain also metal ions of e.g. Cu, Fe, and Zn, and such ions are known to interact with Aβ peptides and modulate their aggregation and toxicity. The interactions between Aβ peptides and Li^+^ ions have however not been well investigated. Here, we use a range of biophysical techniques to characterize *in vitro* interactions between Aβ peptides and Li^+^ ions. We show that Li^+^ ions display weak and non-specific interactions with Aβ peptides, and have minor effects on Aβ aggregation. These results indicate that possible beneficial effects of Li on AD pathology are not likely caused by direct interactions between Aβ peptides and Li^+^ ions.

## INTRODUCTION

Alzheimer’s disease (AD) is still an incurable disease and the main cause of age-related dementia worldwide (Querfurth & LaFerla, 2010; Prince *et al*., 2015; Frozza *et al*., 2018), despite decades of research on putative drugs (Luo *et al*., 2013; Wärmländer *et al*., 2013; Decker & Munoz-Torrero, 2016; Kisby *et al*., 2019). In addition to signs of neuroinflammation and oxidative stress (Agostinho *et al*., 2010; Al-Hilaly *et al*., 2013; Wang *et al*., 2014; Heppner *et al*., 2015; Regen *et al*., 2017), AD brains display characteristic lesions in the form of intracellular neurofibrillary tangles, consisting of aggregated hyperphosphorylated tau proteins (Goedert, 2018; Gibbons *et al*., 2019), and extracellular amyloid plaques, consisting mainly of insoluble fibrillar aggregates of amyloid-β (Aβ) peptides (Glenner & Wong, 1984; Querfurth & LaFerla, 2010). These Aβ fibrils and plaques are the end-product of an aggregation process (Querfurth & LaFerla, 2010; Luo *et al*., 2016; Selkoe & Hardy, 2016) that involves extra- and/or intracellular formation of intermediate, soluble, and likely neurotoxic Aβ oligomers (Luo *et al*., 2014; Selkoe & Hardy, 2016; Sengupta *et al*., 2016; Lee *et al*., 2017) that can spread from neuron to neuron via exosomes (Nath *et al*., 2012; Sardar Sinha *et al*., 2018).

The Aβ peptides comprise 37-43 residues and are intrinsically disordered in aqueous solution. They have limited solubility in water due to the hydrophobicity of the central and C-terminal Aβ segments, which may fold into a hairpin conformation upon aggregation (Abelein *et al*., 2014; Baronio *et al*., 2019). The charged N-terminal segment is hydrophilic and readily interacts with cationic molecules and metal ions (Luo *et al*., 2013; Luo *et al*., 2014; Tiiman *et al*., 2016; Wallin *et al*., 2016; Wallin *et al*., 2017; Owen *et al*., 2019; Wallin *et al*., 2020), while the hydrophobic C-terminal segment can interact with membranes where Aβ may exert its toxicity (Österlund *et al*., 2018; Wärmländer *et al*., 2019). The interactions between Aβ and metal ions are of particular interest (Duce *et al*., 2011; Wärmländer *et al*., 2013; Mital *et al*., 2015; Wärmländer *et al*., 2019; Wallin *et al*., 2020), as altered metal concentrations indicative of metal dyshomeostasis are a prominent feature in the brains and fluids of AD patients (Wang *et al*., 2015; Szabo *et al*., 2016), and because AD plaques contain elevated amounts of metal ions of e.g. Cu, Fe, and Zn (Beauchemin & Kisilevsky, 1998; Lovell *et al*., 1998; Miller *et al*., 2006).

Interestingly, although the role of metal ions in AD pathogenesis remains debated (Duce *et al*., 2011; Modgil *et al*., 2014; Chin-Chan *et al*., 2015; Mital *et al*., 2015; Adlard & Bush, 2018; Huat *et al*., 2019; Wärmländer *et al*., 2019), monovalent ions of the alkali metal lithium [i.e., Li^+^ ions] may provide beneficial effects to patients with neurodegenerative disorders such as amyotrophic lateral sclerosis (ALS) (Fornai *et al*., 2008; Morrison *et al*., 2013) or AD (Engel *et al*., 2008; Mauer *et al*., 2014; Sutherland & Duthie, 2015; Decker & Munoz-Torrero, 2016; Donix & Bauer, 2016; Morris & Berk, 2016; Kerr *et al*., 2018; Hampel *et al*., 2019; Kisby *et al*., 2019; Priebe & Kanzawa, 2020). Lithium salts are commonly used in psychiatric medication, even though it is not understood how the Li^+^ ions affect the molecular mechanisms underlying the psychiatric disorders (Dell’Osso *et al*., 2016). Unlike other pharmaceuticals, Li^+^ is widely non-selective in its biochemical effects, possibly due to its general propensity to inhibit the many enzymes that have magnesium as a cofactor (Ge & Jakobsson, 2018).

Cell and animal studies have provided clues regarding how Li^+^ ions may affect the AD disease pathology (Nery *et al*., 2014; Sofola-Adesakin *et al*., 2014; Zhao *et al*., 2014; Budni *et al*., 2017; Habib *et al*., 2017; Cardillo *et al*., 2018; Kerr *et al*., 2018; Habib *et al*., 2019; Rocha *et al*., 2020; Wilson *et al*., 2020). Due to its ability to down-regulate translation, Li^+^ caused a reduction in protein synthesis and thus Aβ_42_ levels in an adult-onset Drosophila model of AD (Sofola-Adesakin *et al*., 2014). Li^+^ reduces Aβ production by affecting the processing/cleavage of the amyloid-β precursor protein (AβPP) in cells and mice, presumably by down-regulating the levels of phosphorylated APP. A main target of Li^+^ is the glycogen synthase kinase 3-beta (GSK-3β) (Ryves & Harwood, 2001) which is implicated in AD pathogenesis (Caccamo *et al*., 2007; Forlenza *et al*., 2014). In AβPP-transgenic mice, reduced activation of the GSK-3β enzyme was associated with decreased levels of APP phosphorylation that resulted in decreased Aβ production (Rockenstein *et al*., 2007). One study on mice with traumatic brain injury reported that Li^+^-treatment improved spatial learning and reduced Aβ production, possibly by reducing the levels of both AβPP and the AβPP-cleaving enzyme BACE1 (Yu *et al*., 2012). More recent mice studies have reported that treatment with Li^+^ ions improved Aβ clearance from the brain (Pan *et al*., 2018), reduced oxidative stress levels (Xiang *et al*., 2020), improved spatial memory (Habib *et al*., 2019), and reduced the amounts of Aβ plaques and phosphorylated tau while also improving spatial memory (Liu *et al*., 2020).

Only a few studies have however investigated how Li^+^ ions could affect the molecular events that appear to underlie AD pathology, such as Aβ aggregation. One study showed that increased ionic strength, i.e. 150 mM of NaF, NaCl, or LiCl, significantly accelerated the kinetics of Aβ amyloid formation, by promoting surface-catalyzed secondary nucleation reactions (Abelein *et al*., 2016). Another recent study used molecular dynamics simulations to find small but distinct differences in how the three monovalent Li^+^, N^+^, and K^+^ ions interact with Aβ oligomers (Huraskin & Horn, 2019). The therapeutic effect of Li^+^ on Aβ plaque quality and toxicity has been reported in mice, where Li^+^ treatment before pathology onset induced smaller plaques with higher Aβ compaction, reduced oligomeric-positive halo, and attenuated capacity to induce neuronal damage (Trujillo-Estrada *et al*., 2013). One hypothesis is that these neuroprotective effects of Li^+^ could be mediated by modifications of the plaque toxicity through the astrocytic release of heat shock proteins (Trujillo-Estrada *et al*., 2013).

Here, we use a range of biophysical techniques to characterize the *in vitro* interactions between Li^+^ ions and Aβ peptides, and how such interactions affect the Aβ amyloid aggregation processes and fibril formation.

## MATERIALS AND METHODS

### Sample preparation

Recombinant Aβ_40_ peptides were purchased from AlexoTech AB (Umeå, Sweden) in either unlabeled or uniformly ^15^N-labeled form. The lyophilized peptides were stored at −80 °C. Samples were dissolved to monomeric form immediately before each measurement. The peptides were first dissolved in 10 mM NaOH, and then sonicated in an ice-bath to avoid having pre-formed aggregates in the sample solutions. Next, the samples were diluted in 20 mM buffer of either sodium phosphate or MES (2-[N-morpholino]ethanesulfonic acid). All preparation steps were performed on a bed of ice, and the peptide concentration was determined by weight. LiCl salt was purchased from Merck & Co. Inc. (USA), and MES hydrate was purchased from Sigma-Aldrich (USA).

Synthetic Aβ_42_ peptides were purchased from JPT Peptide Technologies (Germany) and used to prepare monomeric solutions via size exclusion chromatography. 1 mg of lyophilized Aβ_42_ powder was dissolved in 250 mL DMSO. A Sephadex G-250 HiTrap desalting column (GE Healthcare, Uppsala) was equilibrated with 5 mM NaOH solution (pH=12.3), and washed with 10-15 mL of 5 mM NaOD, pD=12.7 (Glasoe & Long, 1960) solution. The peptide solution in DMSO was applied to the column, followed by injection of 1.25 mL of 5 mM NaOD. Collection of peptide fractions in 5 mM NaOD on ice was started at a 1 mg/mL flow rate. Ten fractions of 1 mL volumes were collected in 1.5 mL Eppendorf tubes. The absorbance for each fraction at 280 nm was measured with a NanoDrop instrument (Eppendorf, Germany), and peptide concentrations were determined using a molar extinction coefficient of 1280 M^-1^cm^-1^ for the single Tyr in Aβ_42_ (Edelhoch, 1967). The peptide fractions were flash-frozen in liquid nitrogen, covered with argon gas on top in 1.5 mL Eppendorf tubes, and stored at −80°C until used. Sodium dodecyl sulfate (SDS)-stabilized Aβ_42_ oligomers of two well-defined sizes (approximately tetramers and dodecamers) were prepared according to a previously published protocol (Barghorn *et al*., 2005), but in D_2_O, at 4-fold lower peptide concentration and without the original dilution step (Vosough & Barth, manuscript). The reaction mixtures (100 μM Aβ_42_ in PBS and containing 0.05 % or 0.2% SDS) were incubated together with 0-10 mM LiCl at 37 °C for 24 hours, and then flash-frozen in liquid nitrogen and stored at −20°C for later analysis.

### Thioflavin T kinetics

A FLUOstar Omega microplate reader (BMG LABTECH, Germany) was used to monitor the effect of Li^+^ ions on Aβ aggregation kinetics, 20 μM monomeric Aβ_40_ peptides were incubated in 20 mM MES buffer, pH 7.35, together with different concentrations of LiCl (0, 20 μM, 200 μM, 2000 μM) and 50 μM Thioflavin T (ThT). ThT is a fluorescent benzothiazole dye, and its fluorescence intensity increases when bound to amyloid aggregates (Gade Malmos *et al*., 2017). Samples were placed in a 96-well plate where the sample volume in each well was 100 μL, four replicates per Li^+^ concentration were measured, the temperature was +37 °C, excitation of the ThT dye was at 440 nm, the ThT fluorescence emission at 480 nm was measured every five minutes, each five-minute cycle involved 140 seconds of shaking at 200 rpm, the samples were incubated for a total of 15 hours, and the assay was repeated three times. To derive parameters for the aggregation kinetics, the ThT fluorescence curves were fitted to the sigmoidal equation 1:

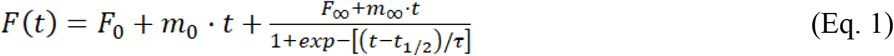

where F_0_ and F_∞_ are the intercepts of the initial and final fluorescence intensity baselines, m_0_ and m_∞_ are the slopes of the initial and final baselines, t_½_ is the time needed to reach halfway through the elongation phase (i.e., aggregation half-time), and τ is the elongation time constant (Gade Malmos *et al*., 2017). The apparent maximum rate constant, r_max_, for the growth of fibrils is given by 1/τ.

### Tyrosine fluorescence quenching

The binding affinity between Aβ_40_ peptides and Li^+^ ions was evaluated from Cu^2+^/ Li^+^ binding competition experiments (Wallin *et al*., 2020). The affinity of the Cu^2+^·Aβ_40_ complex was measured *via* the quenching effect of Cu^2+^ ions on the intrinsic fluorescence of Y10, which is the only fluorophore in native Aβ peptides. The fluorescence emission intensity at 305 nm (excitation wavelength 276 nm) was recorded at 20 °C using a Jobin Yvon Horiba Fluorolog 3 fluorescence spectrophotometer (Longjumeau, France). The titrations were carried out by consecutive additions of 0.8 – 3.2 μL aliquots of either 2, 10, or 50 mM stock solutions of CuCl_2_ to 800 μL of 10 μM Aβ_40_ in 20 mM MES buffer, pH 7.35, in a quartz cuvette with 4 mm path length. After each addition of CuCl_2_ the solution was stirred for 30 seconds before recording fluorescence emission spectra. Copper titrations were conducted for Aβ_40_ samples both in the absence and the presence of 1 mM LiCl. The dissociation constant of the Cu^2+^· Aβ_40_ complex was determined by fitting the Cu^2+^ titration data to equation 2:

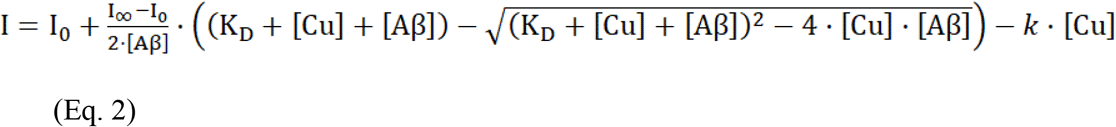

where I_0_ is the initial fluorescence intensity without Cu^2+^ ions, I_∞_ is the steady-state (saturated) intensity at the end of the titration series, [Aβ] is the peptide concentration, [Cu] is the concentration of added Cu^2+^ ions, K_D_ is the dissociation constant of the Cu^2+^· Aβ_40_ complex, and k is a constant accounting for the concentration-dependent quenching effect induced by free (non-bound) Cu^2+^ ions that may collide with the Y10 residue (Lindgren *et al*., 2013). This model assumes a single binding site. As no corrections for buffer conditions are made, i.e. in terms of possible interactions between the metal ions and the buffer, the calculated dissociation constant should be considered to be apparent.

### Atomic force microscopy imaging

Samples of 20 μM Aβ_40_ in 5 mM MES buffer (total volume 100 μL, pH 7.35) with either 0, 20 μM, 200 μM or 2 mM LiCl were put in small Eppendorf tubes and incubated for 72 hours at 37°C under continuous shaking at 300 rpm. A droplet (1 μL) of incubated solution was then placed on a fresh silicon wafer (Siegert Wafer GmbH, Germany) and left to dry for 2 minutes. Next, 10 uL of Milli-Q H_2_O was carefully added to the semi-dried sample droplet and soaked immediately with a lint-free wipe, to remove excess salts in a mild manner. The wafer was left to dry in a covered container to protect it from dust, and atomic force microscopy (AFM) images were recorded on the same day. A neaSNOM scattering-type near-field optical instrument (Neaspec GmbH, Germany) was used to collect the AFM images under tapping mode (Ω: 280 kHz, tapping amplitude 50-55 nm) using Pt/Ir-coated monolithic ARROW-NCPt Si tips (NanoAndMore GmbH, Germany) with tip radius <10 nm. Images were acquired on 2.5 x 2.5 μm scan-areas (200 x 200-pixel size) under optimal scanspeed (i.e., 2.5 ms/pixel). The recorded images were minimally processed using the Gwyddion software where a basic plane leveling was performed (Nečas & Klapetek, 2012).

### Nuclear magnetic resonance spectroscopy

An Avance 700 MHz nuclear magnetic resonance (NMR) spectrometer (Bruker Inc., USA) equipped with a cryoprobe was used to investigate possible interactions between Li^+^ ions and monomeric Aβ_40_ peptides at the atomic level. 2D ^1^H-^15^N-HSQC spectra of 92.4 μM monomeric ^15^N-labeled Aβ_40_ peptides were recorded at 5 °C with 90/10 H_2_O/D_2_O, either in 20 mM MES buffer at pH 7.35 or in 1x PBS buffer (137 mM NaCl, 2.7 mM KCl, and 10 mM phosphate pH 7.4), before and after additions with LiCl. Diffusion measurements were performed on a sample of 55 μM unlabeled monomeric Aβ_40_ peptide in 20 mM sodium phosphate buffer, 100 % D_2_O, pD 7.5, at 5 °C, before and after additions with LiCl dissolved in D_2_O. The diffusion experiments employed pulsed field gradients (PFG:s) according to previously described methods (Danielsson *et al*., 2002), and methyl group signals between 0.7-0.4 ppm were integrated, evaluated, and corrected for the viscosity of D_2_O at 5 °C (Cho *et al*., 1999). All NMR data was processed with the Topspin version 3.6.2 software, and the HSQC crosspeak assignment for Aβ_40_ in buffer is known from previous studies (Danielsson *et al.*, 2006).

### Circular dichroism spectroscopy

Circular dichroism (CD) spectra of 20 μM Aβ_40_ peptides in 20 mM sodium phosphate buffer, pH 7.35, were recorded at 20 °C using a Chirascan CD spectrometer (Applied Photophysics, UK) and a quartz cuvette with an optical path length of 2 mm. Measurements were done between 190 – 250 nm, with a step size of 1 nm and a sampling time of 4 s per data point. First, a spectrum was recorded for Aβ_40_ alone. Next, micelles of 50 mM SDS were added to create a membrane-mimicking environment. Finally, LiCl was titrated to the sample in steps up to a concentration of 512 μM.

### Blue native polyacrylamide gel electrophoresis

Homogeneous solutions of 100 μM Aβ_42_ oligomers prepared in presence and absence of 0 – 10 mM Li^+^ ions were analyzed with blue native polyacrylamide gel electrophoresis (BN-PAGE) using the Invitrogen system. 4-16% Bis-Tris Novex gels (ThermoFisher Scientific, USA) were loaded with 10 μL of Aβ_42_ oligomer samples alongside the Amersham High Molecular Weight calibration kit for native electrophoresis (GE Healthcare, USA). The gels were run at 4 °C using the electrophoresis system according to the Invitrogen instructions (ThermoFisher Scientific, USA), and then stained using the Pierce Silver Staining kit according to the instructions (ThermoFisher Scientific, USA).

### Infrared spectroscopy

Fourier-transformed infrared (FTIR) spectra of Aβ_42_ oligomers were recorded in transmission mode on a Tensor 37 FTIR spectrometer (Bruker Optics, Germany) equipped with a sample shutter and a liquid nitrogen-cooled MCT detector. The unit was continuously purged with dry air during the measurements. 8-10 μL of the 80 μM Aβ_42_ oligomer samples, containing 0 – 10 mM LiCl, were put between two flat CaF2 discs separated by a 50 μm plastic spacer covered with vacuum grease at the periphery. The assembled discs were mounted in a holder inside the instrument’s sample chamber. The samples were allowed to sit for at least 15 minutes after closing the chamber lid, to avoid interference from CO_2_ and H_2_O vapor. FTIR spectra were recorded at room temperature in the 1900-800 cm^-1^ range, with 300 scans for both background and sample spectra, using a 6 mm aperture and a resolution of 2 cm^-1^. The light intensities above 2200 cm^-1^ and below 1500 cm^-1^ were blocked with respectively a germanium filter and a cellulose membrane (Baldassarre & Barth, 2014). The spectra were analyzed and plotted with the OPUS 5.5 software, and second derivatives were calculated with a 17 cm^-1^ smoothing range.

## RESULTS

### ThT fluorescence: influence of Li^+^ ions on Aβ_40_ aggregation

The fluorescence intensity of the amyloid-marker molecule ThT was measured when 20 μM Aβ_40_ samples were incubated for 15 hours together with different concentrations of LiCl (Fig. 1). Fitting Eq. 1 to the ThT fluorescence curves yielded the kinetic parameters t1/2 (aggregation half-time) and r_max_ (maximum aggregation rate) (Fig. 1; Table 1). For 20 μM Aβ_40_ alone, the aggregation half-time is approximately 3.7 hours under the experimental conditions used, and the maximum aggregation rate is 0.5 hours^-1^ (Table 1). These kinetic parameters are not much affected by addition of LiCl in 1:1 or 10:1 Li^+^:Aβ ratios. At the Li^+^:Aβ ratio of 100:1, the r_max_ value remains largely unaffected while the aggregation halftime is increased to almost 5 hours (Fig 1; Table 1). The observation that a Li^+^:Aβ ratio of 100:1 is required to shift the ThT curve clearly shows that Li^+^ ions do not have a strong effect on the Aβ_40_ aggregation kinetics.

**Fig. 1.**
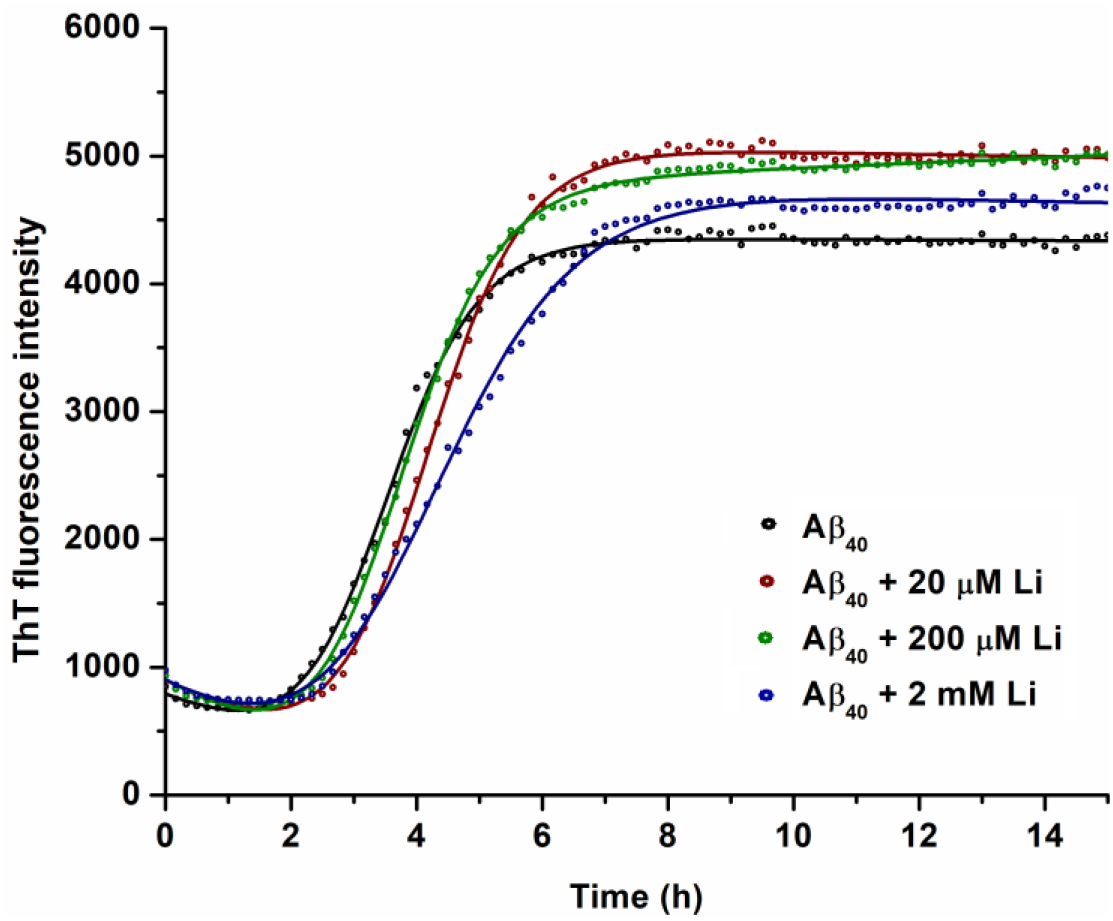
Amyloid fibril formation monitored by ThT aggregation. Samples of 20 μM Aβ_40_ peptides in 20 mM MES buffer, pH 7.35, were incubated at +37 °C together with 50 μM Thioflavin-T and different concentrations of LiCl: 0 μM – black; 20 μM – red; 200 μM – green; 2000 μM – blue. The circles represent average data points for four replicates, while the solid lines are derived from fitting to Eq. 1.

**Table 1.** Kinetic parameters for Aβ_40_ fibril formation, i.e. aggregation half-time (t_1/2_) and maximum aggregation rate (r_max_), derived from fitting the curves in Fig. 1 to Eq. 1.

| | 20 $\mu\text{M}$ $\text{A}\beta_{40}$ | 1:1 $\text{Li}^+:\text{A}\beta$ | 10:1 $\text{Li}^+:\text{A}\beta$ | 100:1 $\text{Li}^+:\text{A}\beta$ |
| --- | --- | --- | --- | --- |
| $t_{1/2}$ [hours] | $3.7 \pm 0.7$ | $3.6 \pm 2.1$ | $3.8 \pm 0.6$ | $4.9 \pm 1.3$ |
| $r_{\text{max}}$ [$\text{hours}^{-1}$] | $0.5 \pm 0.1$ | $0.4 \pm 0.2$ | $0.5 \pm 0.2$ | $0.4 \pm 0.1$ |

### AFM imaging: effects of Li^+^ ions on the morphology of Aβ_40_ aggregates

AFM images (Fig. 2) were recorded for the aggregation products of 20 μM Aβ_40_ peptide, incubated for three days without or with LiCl. The control sample without Li^+^ displays long (> 2 μm) amyloid fibrils that are around 6 nm thick, together with small (< 2 nm) aggregate particles that may be protofibrils (Fig. 2A). The distribution and sizes of these aggregates are rather typical for Aβ_40_ aggregates formed *in vitro* (Luo *et al*., 2014). The Aβ_40_ samples incubated in the presence of different concentrations of Li^+^ ions display amyloid fibrils of similar size and shape, although these fibrils are more densely packed and they appear to be more numerous (Figs. 1B, 1C, 1D). Compared to the control sample, there are fewer small (< 2 nm) aggregate particles in the samples incubated together with Li^+^ ions in 10:1 and 100:1 Li^+^: Aβ ratios. This suggests that Li^+^ ions may induce some differences in the Aβ_40_ aggregation process.

**Fig. 2.**
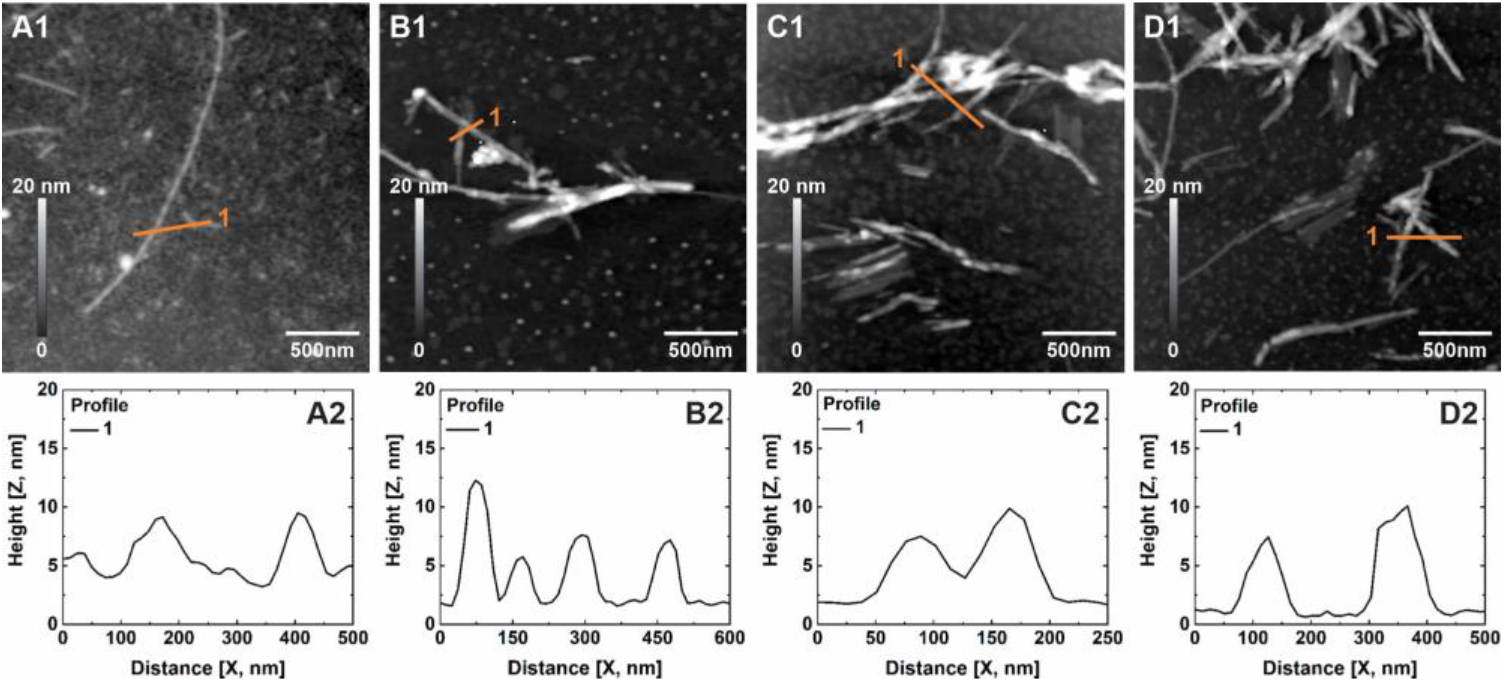
Solid state AFM images (A1-D1) of aggregates of 20 μM Aβ_40_, incubated in 5 mM MES buffer, pH 7.35, for 72 hours at +37 °C with 300 rpm shaking, together with different concentrations of LiCl. A. control sample - no LiCl; B. 20 μM LiCl; C. 200 μM LiCl; D. 2 mM LiCl. The height profile graphs (A2-D2) below the AFM images correspond to the crosssections of Aβ_40_ fibrils shown as white lines in the AFM images.

### NMR spectroscopy: interactions between Li^+^ ions and Aβ_40_ monomers

High-resolution liquid phase NMR experiments were conducted to investigate if residue-specific molecular interactions could be observed between Li^+^ ions and monomeric Aβ_40_ peptides. 2D ^1^H-^15^N-HSQC spectra showing the amide crosspeak region for 92.4 μM monomeric ^15^N-labeled Aβ_40_ peptides are presented in Fig. 3A, before and after addition of LiCl in 1:1, 1:10, and 1:100 Aβ:Li^+^ ratios in 20 mM MES buffer, 7.35. Addition of Li^+^ ions induces loss of signal intensity mainly for amide crosspeaks corresponding to residues in the N-terminal half of the peptide, indicating selective Li^+^ interactions in this region (Fig. 3B). The effects are clearly concentration-dependent. Because Li^+^ ions are not paramagnetic, this loss of signal intensity is arguably caused by chemical exchange related to structural rearrangements induced by the Li^+^ ions. As no chemical shift changes are observed for the crosspeak position (Fig. 3A), these Li^+^-induced secondary structures appear to be short-lived. Figs. 3C and 3D show the results of similar experiments carried out in 1x PBS buffer, i.e. 137 mM NaCl, 2.7 mM KCl, and 10 mM phosphate pH 7.4. Here, the Li^+^ ions induce virtually no changes in the crosspeak intensities, showing that the weak Li^+^/Aβ_40_ interactions observed in pure MES buffer (Figs. 3A,B) disappear when the buffer and ionic strength correspond to physiological conditions.

**Fig. 3.**
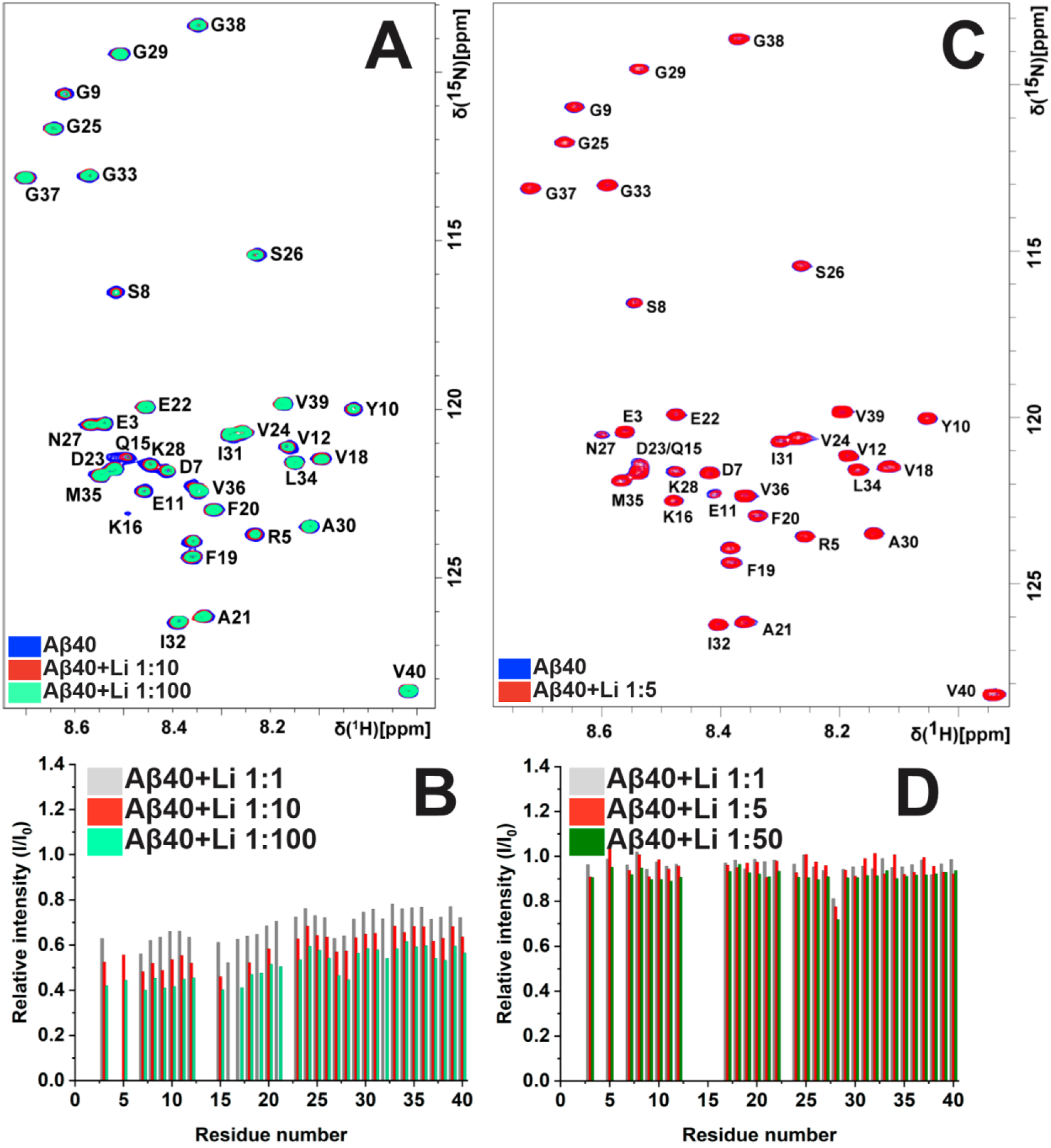
NMR experiments for interactions between Aβ_40_ monomers and Li^+^ ions. (A) 2D ^1^H- ^15^N-HSQC spectra of 92.4 μM ^15^N-labeled Aβ_40_ peptides in 20 mM MES buffer, pH 7.35 at +5 °C, recorded for Aβ_40_ peptides alone (dark sky blue) and in the presence of either 924 μM LiCl (1:10 Aβ:Li ratio; passion red) or 9.24 mM LiCl (1:100 Aβ:Li ratio; Robin egg blue). (B) Relative intensities of Aβ_40_ residue crosspeaks shown in (A), after addition of LiCl in 1:1, 1:10, and 1:100 Aβ:Li ratios. (C and D) similar experiments as in A and B, but carried out in the presence of 1x PBS buffer, and for Aβ:Li ratios of 1:1, 1:5, and 1:50.

Diffusion measurements were carried out for 55 μM Aβ_40_ peptides in D_2_O, before and after addition of LiCl in 1:1, 20:1, and 100:1 Li^+^:Aβ ratios. Addition of 1:1 Li^+^ produces an increase in the Aβ_40_ diffusion rate by around 4%, i.e. from 5.97·10^-11^ m^2^/s to 6.23·10^-11^ m^2^/s (Figs. 4A and 4B). This somewhat faster diffusion is likely caused by the Aβ_40_ peptide adopting a slightly more compact structure in the presence of Li^+^ ions, an effect similar to that previously reported for zinc ions (Abelein *et al*., 2015). Addition of even higher Li^+^ concentrations −20 and 100 times the Aβ_40_ concentration – produces diffusion rates that are similar but a little bit lower than the diffusion rate measured for 1:1 Li^+^:Aβ_40_ ratio, i.e. respectively 6.19·10^-11^ m^2^/s and 6.15·10^-11^ m^2^/s (Figs. 4C and 4D), indicating that the effect of Li^+^ on the Aβ_40_ secondary structure and diffusion has been saturated.

**Fig. 4.**
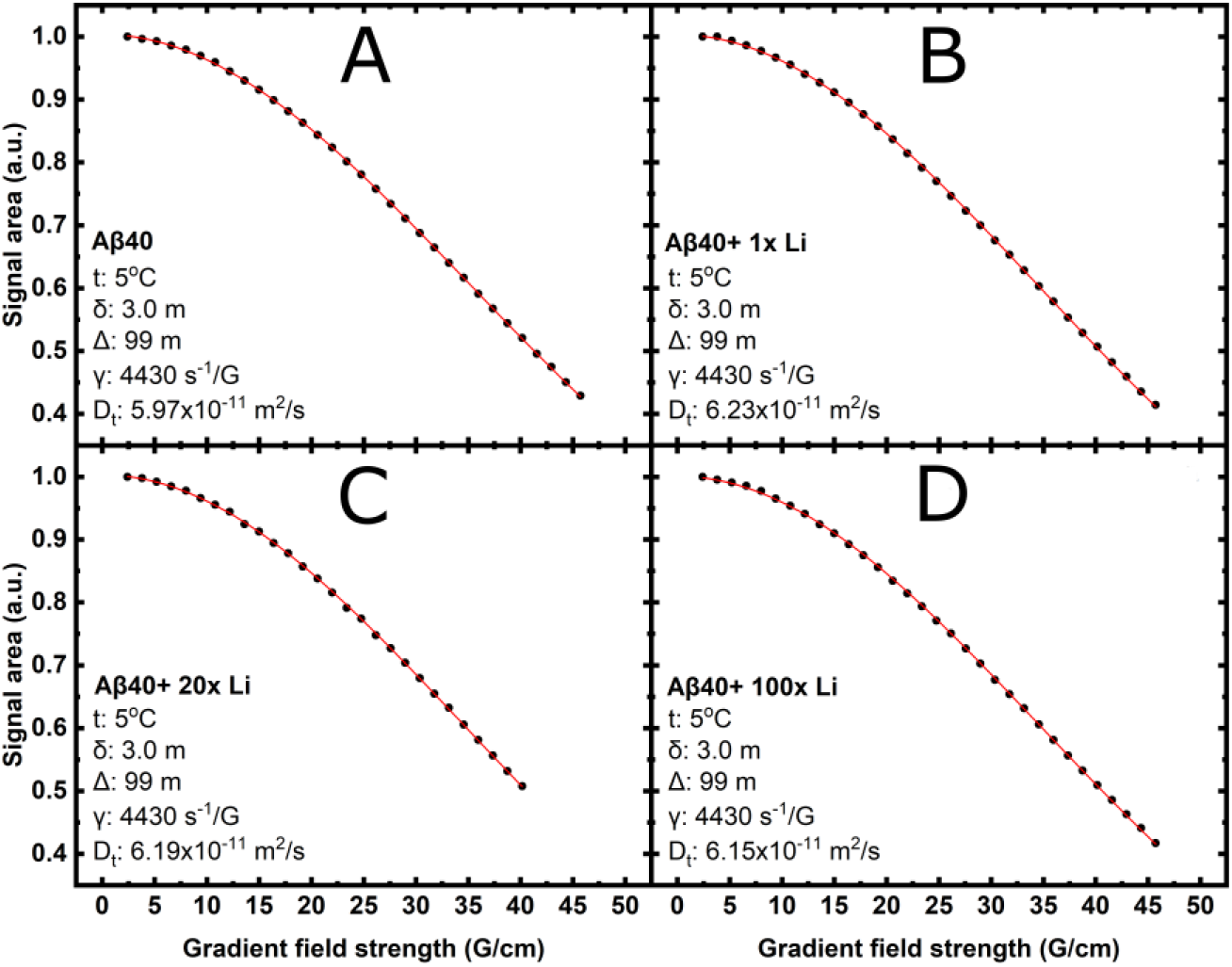
NMR diffusion data for 55 μM Aβ_40_ peptides in sodium phosphate buffer, pH 7.35 at +5 °C, recorded both in absence (A) and presence of different Li^+^ concentrations, i.e. 55 μM (B), 1.1 mM (C), and 5.5 mM (D).

### Fluorescence spectroscopy: Li^+^ binding affinity to the Aβ_40_ monomer

Binding affinities for metal ions to Aβ peptides can often be measured *via* the quenching effect on the intrinsic fluorescence of Y10, the only fluorophore in native Aβ peptides. However, not all metal ions interfere with tyrosine fluorescence, and initial experiments showed that addition of Li^+^ ions does not affect the Aβ_40_ fluorescence. The binding affinity of Li^+^ ions to Aβ_40_ was therefore evaluated from binding competition experiments with Cu^2+^ ions (Danielsson *et al*., 2007; Wallin *et al*., 2020), which induce much stronger tyrosine fluorescence quenching when bound to the peptide than when free in the solution (Lindgren *et al*., 2013). Fig. 5 shows the results of titrating CuCl_2_ to Aβ_40_, both in the absence (red circles) and in the presence (blue triangles) of 1 mM LiCl.

**Fig. 5.**
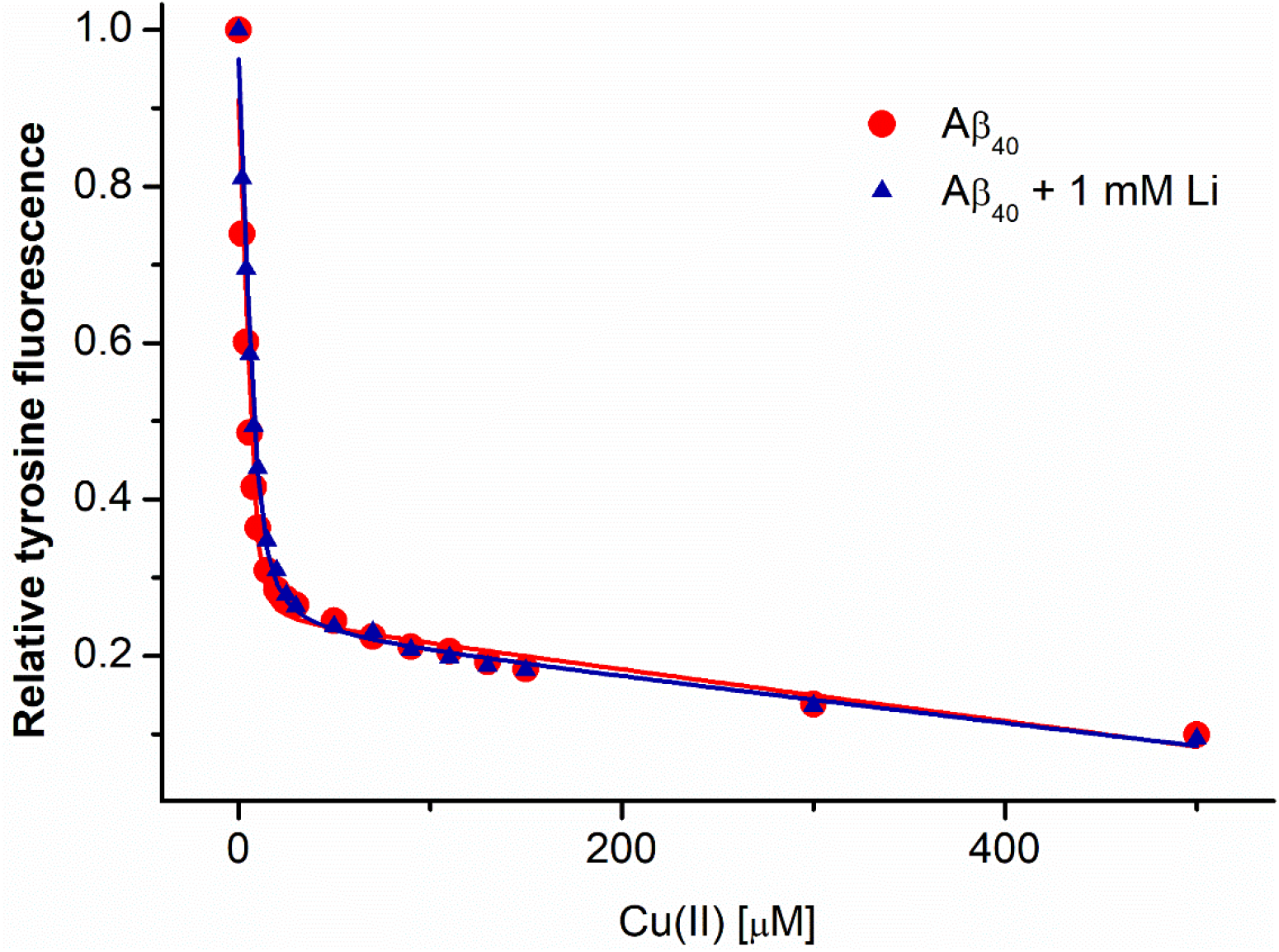
Binding curves for the Cu^2+^·Aβ_40_ complex, obtained from the quenching effect of Cu^2+^ ions on the intrinsic fluorescence of Aβ residue Y10. CuCl_2_ was titrated to 10 μM Aβ_40_ in 20 mM MES buffer, pH 7.35 at 20 °C, both in the absence (red dots) and the presence (blue triangles) of 1 mM LiCl.

Three titrations were carried out for each condition, producing apparent K_D_ values for the Cu^2+^·Aβ_40_ complex of respectively 3.1 μM, 2.1 μM, and 5.1 μM without LiCl, i.e. on average 3.4 ± 1.6 μM, and 2.1 μM, 0.9 μM, and 0.8 μM with LiCl present, i.e. on average 1.3 ± 0.8 μM. The obtained values are in line with earlier fluorescence measurements of the Cu^2+^ binding affinity to the Aβ_40_ peptide, although this affinity is known to vary with the pH, the buffer, and other experimental conditions (Ghalebani *et al*., 2012; Alies *et al*., 2013). The difference between the average measured K_D_ values is not significant at the 5% level with a two-tailed t-test, which shows that Li^+^ ions are not able to compete with Cu^2+^ for binding to Aβ. Thus, the Li^+^ binding affinity for Aβ_40_ is likely in the millimolar range, or weaker.

### CD spectroscopy: effects of Li^+^ ions on Aβ_40_ structure in SDS

Although Aβ peptides are generally disordered in aqueous solutions, they adopt an α-helical secondary structure in membranes and membrane-mimicking environments such as SDS micelles (Tiiman *et al*., 2016; Österlund *et al*., 2018). Thus, the CD spectrum for Aβ_40_ in sodium phosphate buffer displays the characteristic minimum for random coil structure at 198 nm (Fig. 6). Addition of 50 mM SDS, which is well above the critical concentration for micelle formation (Österlund *et al*., 2018), induces an alpha-helical structure with characteristic minima around 208 and 222 nm. Titrating LiCl in concentrations up to 512 μM to the Aβ_40_ sample slightly increases the general CD intensity, but does not change the overall spectral shape – the minima remain at their respective positions. The intensity changes are not caused by dilution of the sample during the titration, as the added volumes are very small, and as dilution would not increase but rather decrease the CD intensity. The observed changes in CD intensity therefore suggest a small but distinct binding effect of LiCl ions. This binding effect appears to be much weaker than the structural rearrangements and Aβ coil-coil-interactions previously reported to be induced by Cu^2+^ ions (Tiiman *et al*., 2016).

**Fig. 6.**
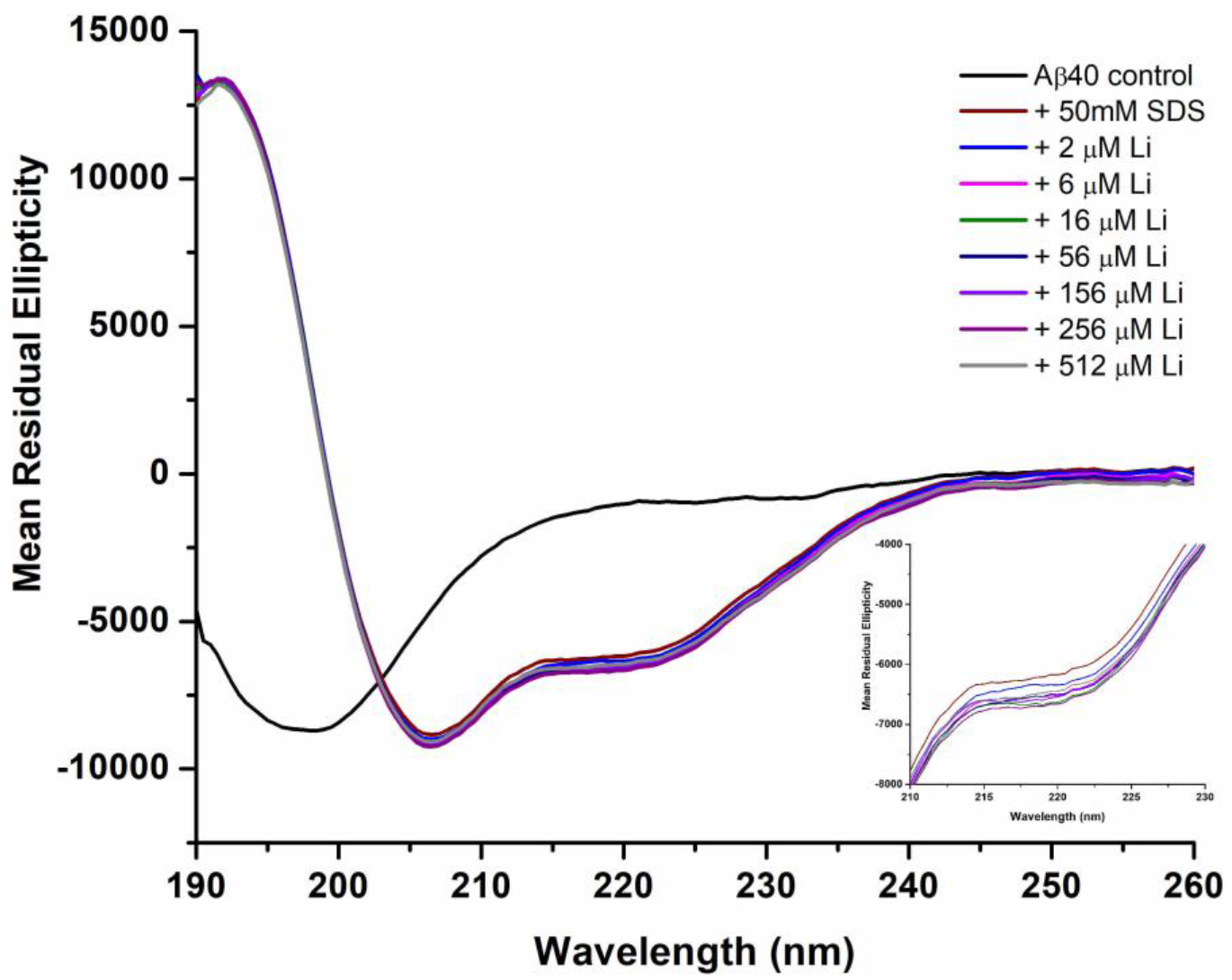
CD spectra of 20 μM Aβ_40_ peptides at 20 °C in 20 mM sodium phosphate buffer, pH 7.35. Spectra were recorded for Aβ in buffer only (black), after addition of 50 mM micellar SDS (brown), and after subsequent addition of between 2 μM (blue) and 512 μM (gray) of LiCl. The inset figure shows a close-up of the CD signals for the LiCl titration in the 210-230 nm range.

### BN-PAGE: effects of Li^+^ ions on Aβ_42_ oligomer formation and stability

Well-defined and SDS-stabilized Aβ_42_ oligomers were prepared in the presence of different amounts of LiCl. SDS treatment of Aβ_42_ peptides at low concentrations (≤ 7 mM) leads to formation of stable and homogeneous Aβ_42_ oligomers of certain sizes and conformations (Barghorn *et al*., 2005; Rangachari *et al*., 2007). As shown in Fig. 7, two sizes of Aβ_42_ oligomers are formed in presence of the two SDS concentrations. In 0.2% (6.9 mM) SDS, small oligomers with a molecular weight (MW) around 16-20 kDa are formed. These oligomers appear to contain a large fraction of tetramers (Vosough & Barth, manuscript). In 0.05% (1.7 mM) SDS, larger oligomers with MWs around 55-60 kD are formed (Barghorn *et al*., 2005). These larger oligomers, which most likely contain twelve Aβ_42_ monomers, display a globular morphology and are therefore sometimes called globulomers (Barghorn *et al*., 2005).

**Fig. 7.**
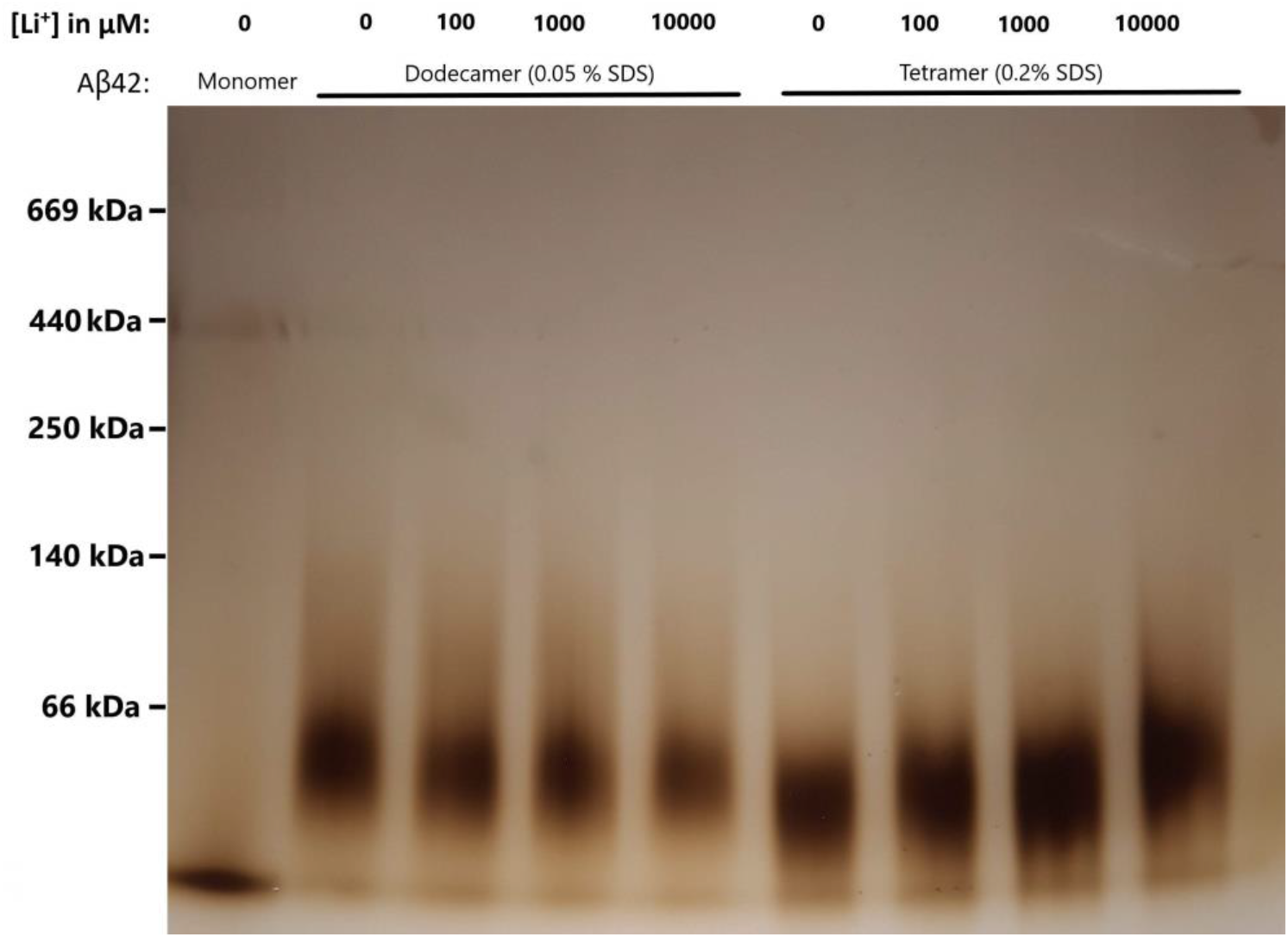
BN-PAGE gel showing the effects of different concentrations of Li^+^ ions on the formation of SDS-stabilized Aβ_42_ oligomers. Lane 1: monomers prepared in 5 mM NaOD. Lanes 2-5: Aβ_42_ globulomers formed after 24 hours of incubation with 0.05% SDS and different LiCl concentrations. Lanes 6-9: Aβ_42_ oligomers formed after 24 hours of incubation with 0.2% SDS and different LiCl concentrations.

All oligomers were analyzed by BN-PAGE instead of by SDS-PAGE to avoid disruption of the non-cross linked Aβ_42_ oligomers by the high (>1%) SDS concentrations used in SDS-PAGE (Bitan *et al*., 2005). As shown in lanes 2-5 and 6-9 of Fig. 7, increasing LiCl concentrations have weak or no effects on the size or homogeneity of the formed Aβ_42_ oligomers, as the bands retain their shape and intensity. Only for the globulomers subjected to the highest LiCl concentration (10 mM) is the intensity of the BN-PAGE band slightly reduced (lane 5, Fig 7).

### FTIR spectroscopy: effects of Li^+^ ions on Aβ_42_ oligomer structure

The secondary structures of Aβ_42_ oligomers formed with different Li^+^ concentrations were studied with FTIR spectroscopy, where the amide I region (1700-1600 cm^-1^) is very sensitive to changes in the protein backbone conformation. The technique is useful also in amyloid research, given its capacity to characterize β-sheets (Barth, 2007; Sarroukh *et al*., 2013).

Fig. 8 shows second derivative IR spectra for the amide I region of Aβ_42_ globulomers (Fig. 8A) and smaller oligomers (Fig. 8B), prepared with different concentrations of Li^+^ ions. Monomeric Aβ_42_ displayed a relatively broad band at 1639-1640 cm^-1^, which is in agreement with the position of the band for disordered (random coil) polypeptides measured in D_2_O (Barth, 2007). For both types of Aβ_42_ oligomers, this main band is much narrower and downshifted by about 10 cm^-1^, while a second smaller band appears around 1685 cm^-1^. This split band pattern is indicative of an anti-parallel β-sheet conformation (Cerf *et al*., 2009).

**Fig. 8.**
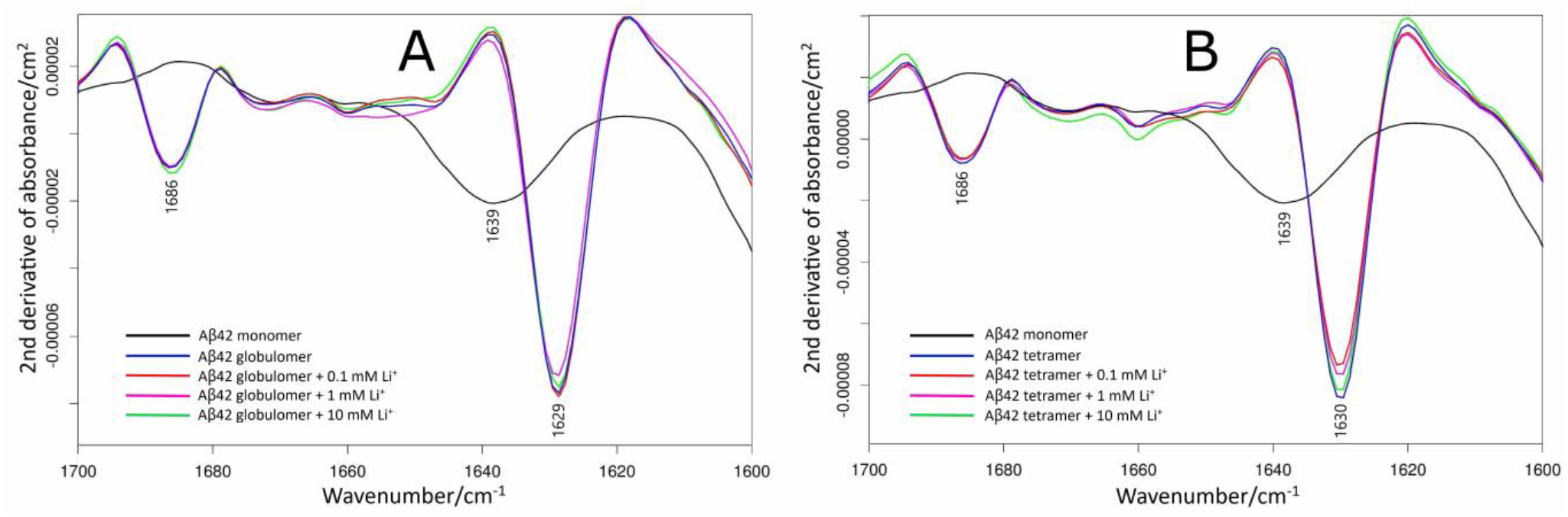
Second derivatives of infrared absorbance spectra for 100 μM Aβ_42_ monomers (black) and 80 μM SDS-stabilized Aβ_42_ oligomers formed in absence (blue) and presence of 0.1 mM (red), 1 mM (purple), and 10 mM (green) of LiCl. The results are shown for Aβ_42_ globulomers at 0.05% SDS (A) and smaller oligomers at 0.2% SDS (B).

Earlier studies in our laboratory have shown a relationship between the position and width of this main band, and the size and homogeneity of the Aβ_42_ oligomers (Vosough & Barth, manuscript). The lower band position of the larger oligomers is in line with this relationship and our previous results, and confirms the different sizes of the oligomers produced at the two SDS concentrations. We have recently observed that a number of transition metal ions induce significant effects on the main band position for Aβ_42_ oligomers (manuscript in preparation). Because the spectra for Aβ_42_ oligomers formed with different amounts of LiCl generally superimpose on the IR spectra for the Li^+^-free oligomers, with no shifts observed for the main band, it appears that Li^+^ ions have no significant effect on the oligomers’ size or secondary structure.

## DISCUSSION

### Lithium as a therapeutic agent

Lithium has no known biological functions in the human body. Li^+^ ions readily pass biological membranes, and are evenly distributed in tissues and easily eliminated via the kidneys (Nordberg *et al*., 2015). Li^+^ ions are however far from inert, and several well-defined medical conditions related to abnormal Li^+^ concentrations exist. In low blood concentrations, Li^+^ is used as a medication for bipolar and schizoaffective disorders (Machado-Vieira *et al*., 2009), but at higher concentrations Li^+^ ions are neurotoxic (Sellers *et al*., 1982; Emilien & Maloteaux, 1996; Nordberg *et al*., 2015; Wen *et al*., 2019). This leaves a narrow therapeutic window of 0.6 −1.2 mM that has to be closely monitored in order to prevent Li^+^ intoxication, which is easily recognized by EEG (Mignarri *et al*., 2013) and treatable by reducing the therapeutic dose. Li^+^ intoxication (>1.5 mM) presents as apathy, vertigo, tremor and gastrointestinal symptoms, in more severe cases confusion, psychosis, myoclonus and cardiac arrhythmias (Nordberg *et al*., 2015). Li^+^ intoxication affects also the kidneys with polyuria and elevated U-albumin although overt renal failure is rare (Nordberg *et al*., 2015).

Treatment of bipolar and schizoaffective disorders with Li^+^ has generated some knowledge about Li^+^ metabolism in the human body (Wen *et al*., 2019; Medic *et al*., 2020). Li^+^ accumulates to some extent in bone (Birch, 1974), and chronic Li^+^ effects are implicated in osteomalacia and severe osteoporosis (Roos, 2014). Patients treated with Li^+^ also show an increased frequency of hypothyroidism and goitre, and widespread effects on several facets of the endocrine system have been noted (Salata & Klein, 1987). The negative effects of Li^+^ on thyroid function have been clearly demonstrated in a study on populations in the Andean Mountains, where natural exposure to Li^+^ is high, and where urinary Li^+^ was found to correlate negatively with free thyroxine (T4) but correlate positively with the pituitary gland hormone thyrotropin (Broberg *et al*., 2011). The toxicity of Li^+^ is further emphasized by studies from regions with naturally elevated concentrations of Li^+^ in potable water, where reduced fetal size has been noted to correlate linearly with increases in blood Li^+^ (Harari *et al.*, 2015).

To what extent Li^+^ treatment reduces the development of AD symptoms is unclear (Engel *et al*., 2008; Mauer *et al*., 2014; Nordberg *et al*., 2015; Sutherland & Duthie, 2015). Bipolar disorder increases the risk of AD when compared to the general population, and Li^+^ treatment seems to reduce this risk (Velosa *et al*., 2020), but the mechanisms mediating this effect are far from elucidated (Kerr *et al*., 2018). In rare cases even regular-dose long-time Li^+^ therapy may cause severe intoxication of the central nervous system, characterized by cerebellar dysfunction and cognitive decline (Emilien & Maloteaux, 1996).

### Lithium interactions with the Aβ_40_ peptide

The NMR (Figs. 3 and 4), fluorescence quenching (Fig. 5), and CD (Fig. 6) experiments show that Li^+^ ions display weak interaction with the Aβ_40_ peptide, where the binding affinity for the Li^+^·Aβ_40_ complex may be in the millimolar range. The IR and CD results show minor or no effects of Li^+^ ions on the secondary structures of Aβ_40_ monomers (Fig. 6) and Aβ_42_ oligomers (Fig. 8). The Li^+^ ions may have a small effect on Aβ aggregation, with minor perturbations on the morphology of aggregated Aβ_40_ fibrils (Fig. 2), and effects on the Aβ_40_ aggregation kinetics (Fig. 1; Table 1) and Aβ_42_ oligomer stability (Fig. 7) only at very high Li^+^ concentrations. These results are in line with previous computer modeling results, which suggest small differences between how the monovalent K^+^, Li^+^, and Na^+^ alkali ions affect Aβ oligomerization (Huraskin & Horn, 2019).

As Aβ_40_ and Aβ_42_ have identical N-terminal sequences, the two peptide variants should interact very similarly with Li^+^ ions, which were found to bind to the N-terminal Aβ region (Fig. 3B). The weak affinity between Aβ_40_ and Li^+^ ions, and the fact that Li^+^ does not efficiently compete with Cu^2+^ ions for Aβ binding (Fig. 4), suggest that Li^+^ ions are not coordinated by specific binding ligands. Instead, Li^+^ likely engages in non-specific electrostatic interactions with the negatively charged Aβ residues, i.e. D1, E3, D7, E11, E22, and D23 (which are located in the N-terminal and central regions).

The weak binding affinity to Aβ_40_ peptides is not caused by Li^+^ ions being monovalent, as e.g. monovalent Ag^+^ ions display rather strong and specific binding to Aβ peptides (Wallin *et al*., 2020). Moreover, divalent Pb^2+^ and trivalent Cr^3+^ ions do not bind strongly to Aβ, while divalent Cu^2+^, Mn^2+^ and Zn^2+^ as well as tetravalent Pb^4+^ ions do (Faller, 2009; Abelein *et al*., 2015; Tiiman *et al*., 2016; Wallin *et al*., 2016; Wallin *et al*., 2017). Thus, Aβ/metal interactions are not governed by the charge of the metal ion, but rather by its specific properties, such as ionic radius and electron configuration (1s^2^ for Li^+^).

It is illustrative to compare the Aβ interactions with Li^+^ ions to the well-studied interactions with Cu^2+^ and Zn^2+^ ions. These two divalent ions display residue-specific interactions with Aβ peptides, displaying binding affinities in the micromolar-nanomolar range and strong effects on Aβ secondary structure, aggregation, and diffusion (Danielsson *et al.*, 2007; Faller, 2009; Lindgren *et al*., 2013; Abelein *et al*., 2015; Tiiman *et al*., 2016; Owen *et al*., 2019). Aβ binding to Cu^2+^ and Zn^2+^ is coordinated mainly by residue-specific interactions with the N-terminal His residues, i.e. H6, H13, and H14 (Faller, 2009; Lindgren *et al*., 2013; Abelein *et al*., 2015; Tiiman *et al*., 2016). The biological relevance of Cu^2+^ and Zn^2+^ ions in AD pathology is demonstrated by their dysregulation in AD patients (Wang *et al.*, 2015; Szabo *et al*., 2016), and by them being accumulated in plaques of Aβ aggregates in AD brains (Beauchemin & Kisilevsky, 1998; Lovell *et al*., 1998; Miller *et al*., 2006). During neuronal signaling Cu^2+^ and Zn^2+^ ions are released into the synaptic clefts (Ayton *et al.*, 2013), where they may interact with Aβ peptides to initiate Aβ aggregation (Branch *et al*., 2017), or modulate the formation and toxicity of Aβ oligomers (Stefaniak & Bal, 2019; Wärmländer *et al*., 2019).

The current results indicate that Li^+^ ions are not able to compete with Cu^2+^ or Zn^2+^ ions for binding to Aβ peptides, and should therefore not be able to influence the *in vivo* effects of Cu^2+^ and Zn^2+^ ions on Aβ aggregation and toxicity. Although high concentrations of Li^+^ showed some effects on Aβ aggregation (Figs. 1–3;7), these effects are likely at least partly related to ionic strength effects (Abelein *et al*., 2016). Under physiological ionic strength, no specific interactions are observed between Aβ_40_ monomers and Li^+^ ions (Fig. 3C,D). Thus, we conclude that the previously reported possible beneficial effects of Li^+^ on Alzheimer’s disease progression (Mauer *et al*., 2014; Sutherland & Duthie, 2015; Kerr *et al*., 2018; Hampel *et al*., 2019; Velosa *et al*., 2020) seem not to be caused by direct interactions between Li^+^ ions and Aβ peptides.

## ACKNOWLEDGMENTS

We thank Elizabeth (Li) Wang for helpful discussions. This work was supported by grants from the Swedish Alzheimer Foundation and the Swedish Research Council to AG, the Swedish Brain Foundation to AG and AB, the Magnus Bergvall Foundation to SW and PR, the Ulla-Carin Lindquist ALS Foundation to PR, and from Olle Engkvist’s Foundation, the Stockholm Region, and Knut and Alice Wallenberg Foundation to AB.

## CONFICT OF INTEREST

The authors declare no conflict of interest.

